# Decoupling environmental suitability from realized recruitment through ontogenetic environmental filtering in a long-lived Mediterranean palm species

**DOI:** 10.64898/2026.08.17.745352

**Authors:** Sebastián Cordero, Florencia R. Pérez, Renata Acuña-Molina, Yerco Contreras-Vera, Trinidad Jorquera-Fonck, Felipe Góngora-Vásquez, Benjamín González-Ramos, Juan Pablo Núñez, Ignacia Roselló, Ágata Sepúlveda-Vásquez, Matías A. Vergara, Francisco E. Fontúrbel

## Abstract

Long-lived plants facing anthropogenic disturbance often exhibit recruitment failure despite persistent reproductive adults, generating extinction debt masked by longevity. However, whether adult presence reliably reflects environmental suitability for recruitment remains unclear. Here, we examine ontogenetic niche differentiation and its consequences for recruitment in *Jubaea chilensis*, an endangered long-lived Mediterranean palm with an aging population. We assigned individuals within the largest known population to four ontogenetic stages and characterized their environmental niches using climatic, edaphic, topographic, and vegetation variables. We then applied spatial and multivariate analyses, including Random Forest models to evaluate environmental segregation and identify predictors of seedling establishment. Age classes occupied significantly different environmental niches, with the greatest differentiation between seedlings and reproductive adults. Saplings and adult differentiation reflected mainly topographic variables at landscape scale, whereas seedling establishment was primarily predicted by microhabitat conditions (vegetation cover heterogeneity, east-facing slope orientation, and soil texture). This pattern is consistent with niche reconfiguring throughout the life cycle, suggesting that adult occurrence and recruitment suitability respond to distinct environmental conditions. Over one-fifth of sampled individuals occupied high-suitability sites without recruitment, suggesting that ontogenetic niche shifts are associated with a spatial decoupling between adult persistence and recruitment, consistent with demographic collapse independent of habitat degradation. This failure is likely mediated by insufficient effective seed dispersal, as the sole disperser (*Octodon degus*) preys on most seeds before dispersal. Conservation strategies based solely on adult distribution may therefore overestimate effective habitat and underestimate extinction risk in long-lived species.

## Introduction

Long-lived species inhabiting ecosystems undergoing anthropogenically driven transformation often experience recruitment limitations that constrain population regeneration (Schweiger et al. 2020). In plant species, this represents a fundamental demographic challenge, as adults can persist for decades or even centuries, but if no younger individuals replace them, populations eventually become extinct (Bialic-Murphy et al. 2022). This phenomenon, known as extinction debt, occurs when adult individuals remain but current conditions no longer support successful recruitment (Kuussaari et al. 2009). Inverted population pyramids, where older adult individuals dominate and saplings and seedlings are scarce, constitute a clear indicator of demographic collapse (Vellend et al. 2006). This pattern is increasingly common in long-lived species worldwide, especially in areas of accelerated climate change or strong anthropogenic disturbance (Allen et al. 2010, Seidl et al. 2017). Despite this problem being widely acknowledged, the underlying mechanisms driving recruitment failure remain largely unknown for most species, particularly whether the presence of adults reliably reflects environmental suitability for recruitment.

Successful plant recruitment implies overcoming several filters operating sequentially and at different spatial scales (Neuschulz et al. 2016). First, plants must overcome biotic filters, where habitat structure modulates animal interactions, leading to positive or negative outcomes (García et al. 2011, Beckman and Rogers 2013). For example, seed fate can be strongly influenced by habitat features, with interacting animal species acting either as seed dispersers or predators (Cordero et al. 2026). However, besides biotic filters, abiotic filters related to climatic, vegetation, edaphic, and topographic conditions of the habitat determine suitable areas for plant establishment (Frei et al. 2018). These filters interact in complex ways, as seeds must first escape animal predation and then reach sites with suitable environmental conditions for establishment and growth (Cordero et al. 2021). Although many studies examine one of these filters, those integrating both biotic and abiotic filters within the same population remain scarce (Švamberková and Lepš 2020). This not only limits our understanding of the causes leading to recruitment failure (Funk 2021), but also of how these interacting filters shape the spatial distribution of successful recruitment relative to adult occurrence.

In addition to biotic and abiotic filters operating simultaneously, plants must also overcome the problem that their ecological niche can differ across life stages. The conditions necessary for seeds to germinate and for seedlings to establish differ from those necessary for adults to survive and reproduce, as each life stage occupies a distinct ecological niche (Bertrand et al. 2011). This implies that the presence of adults is not necessarily indicative of suitable conditions for recruitment, as environmental requirements change throughout ontogeny (Máliš et al. 2016). In long-lived species, this issue becomes critical, as current adult individuals were possibly established in past decades or centuries under quite different environmental conditions (Schweiger et al. 2020). Therefore, adults may persist in sites where recruitment is currently unfeasible under contemporary environmental conditions (Sandel et al. 2025), leading to spatial segregation among age classes (Weng et al. 2017). Understanding this ontogenetic segregation and its underlying environmental drivers is key to predicting where recruitment can take place.

Climate change and anthropogenic disturbance are progressively altering environmental conditions (Pecl et al. 2017), driving the contraction of the regenerative niche of species over time (Donohue et al. 2010, Walck et al. 2011). Mediterranean ecosystems are particularly vulnerable to this phenomenon, as they have experienced aridification processes over the last decades, with a reduction in precipitation and an increase in temperatures (Giorgi and Lionello 2008, Hoerling et al. 2012). Long-lived species with episodic recruitment are especially susceptible to niche contraction, as their recruitment is restricted to specific time frames that can become progressively scarce (Holmgren et al. 2006). Recruitment can therefore fail even at sites occupied by reproductive individuals if the environmental niche required for seedling establishment differs from the niche currently occupied by reproductive adults (Dalgleish et al. 2011). This mismatch generates an extinction debt, whereby populations appear demographically stable despite ongoing and imminent local extinction (Kuussaari et al. 2009).

*Jubaea chilensis* (Arecaceae) represents an ideal study system to examine this phenomenon. *Jubaea chilensis* is a long-lived palm species, reaching ages of over 200 years (Guzmán et al. 2017), which implies that current adults likely established under climatic conditions different from contemporary ones. Previous studies have described sequential biotic filtering during recruitment, accounting for recruitment rates lower than 2% and inverted age structures (Cordero et al. 2021). However, despite biotic filters being well characterized, environmental filters determining where individuals escaping predation can successfully establish remain unknown. Here, we examine whether recruitment failure in *J. chilensis* is explained by environmental filtering across ontogenetic stages and by regenerative niche contraction under recent environmental change. We hypothesize that recruitment is constrained primarily by fine-scale microhabitat conditions, whereas segregation among later ontogenetic stages is driven mainly by landscape-scale variables. Furthermore, we expect current regenerative niche contraction, whereby the conditions occupied by recently established seedlings represent a more restricted subset of those occupied by reproductive individuals established decades ago (∼at least 60 years).

## Methods

### Study area and data collection

During the austral spring of 2025, we conducted field sampling in the Palmar de Ocoa, located within La Campana National Park, which harbors the largest known population of *J. chilensis* (∼70,000 individuals; González et al. 2017). Although legally protected, this site is threatened by multiple anthropogenic pressures, such as fruit harvesting, herbivory by exotic species, wildfires, and seed predation by invasive rodents (Cordero et al. 2021). Nonetheless, it represents the least disturbed of the remaining large *J. chilensis* populations, most of which occur outside protected areas in a heavily fragmented and degraded matrix (Cordero et al. 2021, Cordero et al. 2026). The sites lies in central Chile within the Winter Rainfall-Valdivian Forests Biodiversity Hotspot and is characterized by a Mediterranean climate-type, with precipitations concentrated during the winter and prolonged dry summers (120 mm annual rainfall; Quintanilla and Lozano 2012). Vegetation is dominated by sclerophyllous forest and scrubland formations, where *J. chilensis* is mainly associated with species such as *Quillaja saponaria, Cryptocarya alba, Lithrea caustica*, *Peumus boldus*, *Vachellia caven*, *Retanilla trinervia*, and *Leucostele chiloensis.* The study area exhibits strong vegetation heterogeneity over short distances, including sclerophyllous forests and scrublands and hygrophilous formations that generates contrasting microhabitats, providing sufficient environmental variation to compare habitat associations among age classes (Cordero et al. 2026). We conducted continuous sampling across 56 ha (∼2% of the total population area), georeferencing all individuals and assigning them to age classes representing ontogenetic stages. Four classes were defined based on morphological aspects (González et al. 2017): seedling (entire or weakly pinnate leaves, basal rosette, no stem), saplings (pinnate leaves, rosette borne on a swollen pseudo-stem formed by persistent, compacted petiole bases), non-reproductive adult (visible stem with an elevated crown, no reproductive structures), and reproductive adult (narrowed stem, presence of inflorescences or infructescences).

### Spatial analyses

To assess whether individuals deviate from a random spatial distribution and whether ontogenetic stages exhibit distinct aggregation patterns, we conducted point pattern analyses using the geographic coordinates of sampled individuals. First, we defined an observation window as the minimum convex hull containing all individuals. To characterize the global spatial pattern of the population, we applied Ripley’s K function with edge correction. This analysis evaluates spatial distribution patterns across multiple distance scales (r). K(r) values higher than those expected under randomness indicate spatial aggregation at that scale, whereas lower values suggest regularity or repulsion. Statistical significance was assessed using Monte Carlo simulations, generating 999 random patterns under a Complete Spatial Randomness (CSR) model and constructing 95% confidence envelopes. Then, to evaluate whether aggregation patterns vary ontogenetically, we conducted the Ripley’s K analysis independently for each age class. Additionally, we estimated the Clark-Evans index for each class, which quantifies the ratio between the mean observed nearest-neighbor distance and that expected under spatial randomness. R values < 1 indicate aggregation, R > 1 suggest regularity, and R ≈ 1 indicate randomness. Statistical significance was evaluated using the Clark-Evans Z-test.

Furthermore, to determine whether different age classes tend to occupy spatially segregated zones, we conducted a spatial segregation test on a pattern of marked points, where each individual was tagged according to its age class. This test evaluates whether the spatial distribution of tags (age classes) significantly differs from that expected under a null model generated by random reassignment of tags to the observed positions, using 999 Monte Carlo simulations.

### Environmental niche characterization and differentiation

Given that age classes exhibit spatial segregation, we evaluated whether this pattern reflects differentiation in environmental space. To characterize the environmental niche of *J. chilensis*, we used climatic, vegetational, edaphic, and topographic variables, obtained using the geographic coordinates of each individual. Climatic variables were retrieved from CHELSA v2.1 (∼1 km resolutions; Karger et al. 2017) and comprised Aridity Index (AI), Potential Evapotranspiration (PET), mean annual temperature (bio1), precipitation seasonality (bio15), mean precipitation of the driest quarter (bio17), and mean precipitation of the warmest quarter (bio18). Vegetational variables were derived from NDVI (Normalized Difference Vegetation Index; 30 m resolution) from Landsat for the period 1984-2023. We included mean NDVI (productivity/mean vegetation cover), NDVI CV (coefficient of variation), NDVI seasonal amplitude (mean annual maximum - mean annual minimum), temporal trend NDVI (slope of linear regression between 1984-2023), and ENSO (El Niño-Southern Oscillation) sensitivity, calculated as the correlation between monthly NDVI anomalies (deviations from the 1984-2023 monthly mean) and the Oceanic Niño Index (ONI). On the other hand, edaphic variables were retrieved from SoilGrids v2.0 (250 m resolution; Poggio et al. 2021), including cation exchange capacity, coarse fragments, total nitrogen, organic carbon, pH, water content at field capacity, and content of clay, sand, and silt. Due to the compositional nature of these last three variables, we created a texture index, calculated as log((sand + 1)/(clay + silt + 1)), where positive values indicate coarse textures and negative values fine textures. Lastly, topographic variables were derived from a Digital Elevation Model (DEM) SRTM (30 m resolution), calculating elevation, slope, aspect (transformed into northness = cos(aspect) and eastness = sin(aspect) to avoid the circularity of aspect in degrees), and Topographic Wetness Index (TWI = ln(accumulation area/tan(slope)), which represents the potential for water accumulation across the landscape.

To avoid multicollinearity within each set of environmental variables, we implemented an iterative procedure for variable selection based on Variance Inflation Factor (VIF). We conducted multiple linear regressions for each variable against all remaining variables, estimating VIF and removing those with values >5, then repeating until obtaining a set of variables with values under the defined threshold. All variables were standardized prior to multivariate analyses to prevent variables with larger variances from dominating ordinations. These variables were selected since climate constrains seasonal water availability, soils regulate moisture retention and establishment, vegetation dynamics modulate biotic interactions, and topography shapes microclimatic and hydrological heterogeneity (Mod et al. 2016). Together, these environmental variables influence recruitment and spatial niche dynamics of *J. chilensis* in Mediterranean systems.

To evaluate whether age classes occupy different environmental niches and to identify the variables underlying this differentiation, we applied a Permutational Multivariate Analysis of Variance (PERMANOVA) across all age classes on a matrix of individuals by standardized environmental variables, using Euclidean distances (appropriate for continuous z-scored data), with 9,999 permutations, followed by pairwise comparisons with Bonferroni correction. To visualize the multivariate structure of this differentiation, we performed a non-metric Multidimensional Scaling (nMDS) ordination in two dimensions on the same distance matrix, with 100 random starts, assessing ordination quality using stress values. Environmental variables driving the observed segregation were identified using vector fitting with 999 permutations, retaining variables with R² > 0.3 and p < 0.05.

### Environmental restrictions on recruitment and ontogenetic transitions

To evaluate which environmental variables better predict the presence of *J. chilensis* seedlings (the most critical age class for population viability) and compare these predictors against those differentiating other demographic transitions, we employed a Random Forest approach. We trained a Random Forest model to simultaneously predict the four classes using 1,000 decision trees and mtry = ⌊√18⌋ = 4 (the default √p, rounded down) variables evaluated in each split. Classes imbalance was handled by applying class weights inversely proportional to their frequency. Out-of-Bag (OOB) error and confusion matrix were also calculated to assess their predictive performance. Variable importance was evaluated through Mean Decrease Gini and Mean Decrease Accuracy, with the latter providing more robust results for variables with different scales. Furthermore, we trained a second model to examine how current recruitment is constrained, classifying individuals as seedling and non-seedling. To handle the extreme class imbalance (1:21), we combined class weighting, balanced sampling (n = 15 per class per tree), and 1,000 trees to stabilize model predictions. The model was evaluated using OOB error rate, sensitivity (seedling detection rate), and specificity (non-seedling detection rate). Variable importance was calculated through Mean Decrease Accuracy to identify the most important predictors of seedling presence. Lastly, we trained a third binary model for saplings and reproductive adults to compare environmental filters in different ontogenetic stages. We determined the most important variables by Mean Decrease Accuracy for each model to identify those emerging consistently, which were considered as robust environmental drivers, while specific variables for each model indicate differentiated ontogenetic filters.

### Spatial suitability mapping and niche contraction assessment

First, we predicted spatial suitability for recruitment, identified areas with apparently suitable conditions but lacking regeneration (recruitment failure), and assessed the magnitude of regeneration niche contraction. To this end, we used the binary Random Forest model (seedling vs. non-seedling individuals) described above to predict the probability of being seedling for all 332 sampled individuals. This approach generates an environmental suitability index that quantifies how similar the conditions of each individual are to those under which seedlings currently establish. Predicted probabilities were compared among classes using a Wilcoxon test to verify that observed seedlings exhibit higher predicted suitability than other classes (model validation). A continuous suitability map for the study area was generated using Inverse Distance Weighting (IDW) interpolation with a power parameter of p = 2.0. A regular grid of 100 × 100 points covering the spatial extent of the study area was created, and predicted probabilities from the 332 individuals were interpolated across this grid. To classify individuals and assess recruitment failure, each individual was assigned to one of four categories based on its current condition and predicted suitability: i) observed seedlings (field-recorded seedlings), ii) high suitability without seedlings (non-seedling individuals with predicted probability ≥ 75th percentile), iii) medium suitability (predicted probability between the 25th and 75th percentiles), and iv) low suitability (predicted probability < 25th percentile).

Furthermore, to evaluate whether the regeneration niche has changed between generations, we compared the environmental conditions currently occupied by reproductive adults (“adult niche”) with those occupied by seedlings (“seedling niche”). For the most important variables identified by the binary Random Forest model, we calculated class-specific medians and applied Wilcoxon tests to assess significant differences. The percentage change in the median ((Seedlings - Reproductive adults)/Reproductive adults) ×100 quantified the magnitude and direction of niche change. Finally, we quantified the reduction in regeneration area by calculating the convex hull enclosing individuals of each age class using geographic coordinates. Polygon areas were computed using the Shoelace formula for irregular polygons. The spatial contraction index was defined as IC = 1 - (Area seedlings / Area reproductive adults), where values close to 0 indicate no contraction (seedlings occupy a spatial extent similar to that of adults) and values close to 1 indicate strong contraction (seedlings occupy a much smaller spatial extent).

## Results

### Spatial aggregation and age-class segregation

We observed that the *J. chilensis* population exhibits a statistically significant pattern of spatial aggregation (Fig. 1). Aggregation was evident even at short distances (< 20 m) and remained significant up to distances of approximately 150 m, suggesting that individuals tend to form spatially defined clusters or patches rather than being independently distributed. This pattern varied consistently among age classes. Seedlings showed the most intense spatial aggregation (Clark-Evans R index = 0.23, p < 0.001), indicating that younger individuals are concentrated in highly compact groups. Saplings also showed significant aggregation, although of lower magnitude (R = 0.61, p < 0.001). Non-reproductive adults exhibited a moderate aggregation pattern (R = 0.71, p < 0.001), whereas reproductive adults were also spatially aggregated (R = 0.59, p < 0.001), although less pronounced than in younger classes. The spatial segregation test revealed that age classes are not randomly distributed across space, but instead occupy distinct zones within the study area (p = 0.001).

**Figure 1.**
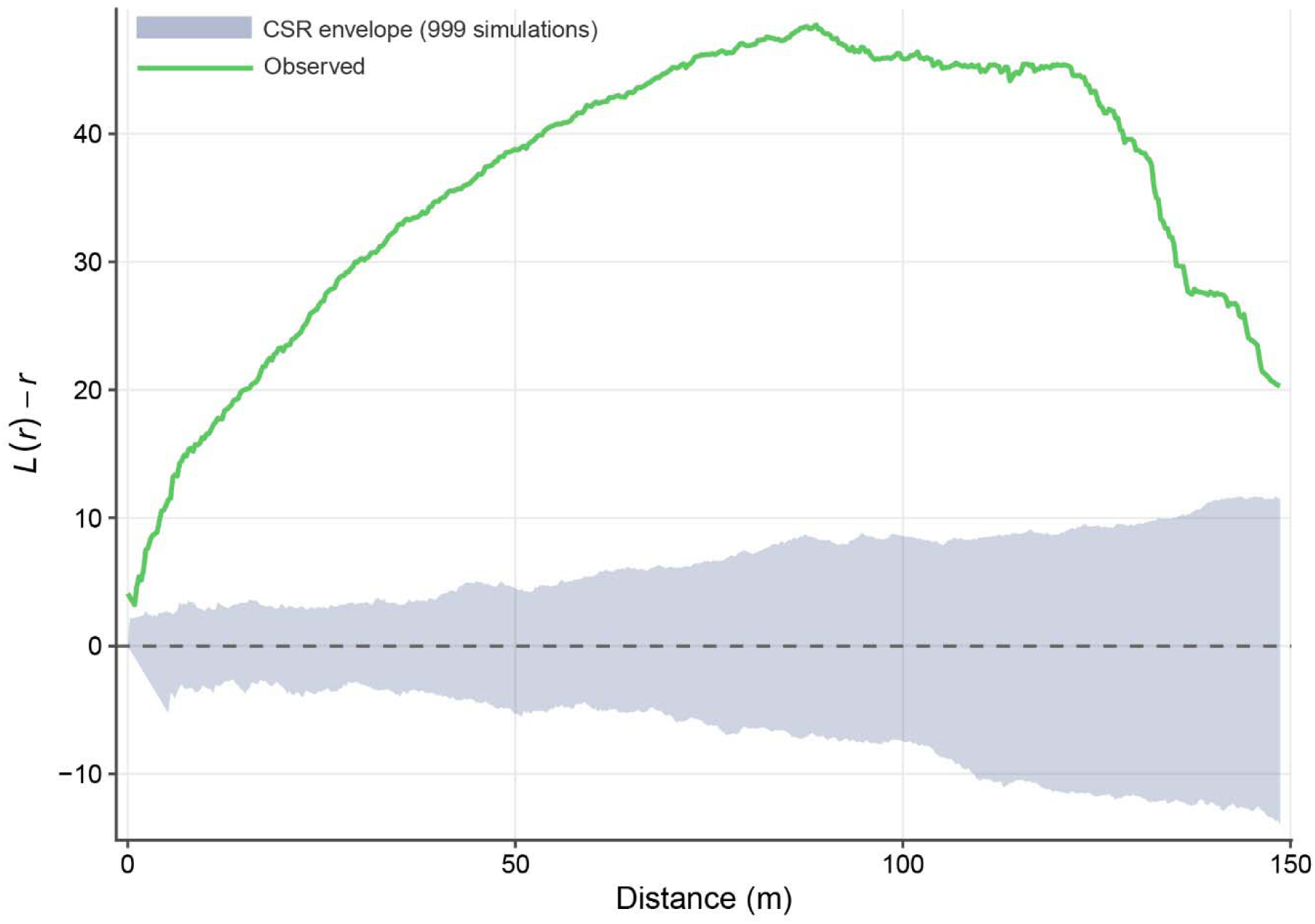
Ripley’s *L*-function analysis of the spatial distribution of *Jubaea chilensis* individuals (n = 332). The x-axis represents spatial scale (distance r, in meters), and the y-axis shows the L(r) − r statistic (values of zero correspond to complete spatial randomness). The green line represents the observed *L*(r) − r values, and the shaded area indicates the simulation envelope under complete spatial randomness (CSR), generated from 999 Monte Carlo simulations. Values above the envelope indicate spatial aggregation, while values within the envelope indicate no significant departure from randomness.

### Age-class segregation in environmental space

Consistent with the observed spatial segregation, age classes occupy significantly different environmental niches (PERMANOVA: F, = 4.51, p < 0.0001), suggesting progressive displacement of environmental conditions between generations. Pairwise comparisons between all class combinations showed that five of six comparisons were statistically significant. Reproductive adults and saplings exhibited the highest differentiation (F = 7.92, R² = 0.03, p_adj_ = 0.001), followed by seedlings and saplings (F = 3.70, R² = 0.06, p_adj_ = 0.01), and reproductive adults and seedlings (F = 3.61, R² = 0.01, p_adj_ = 0.02). The only exception was non-reproductive adults and saplings (p_adj_ = 1.00). Consistent with this statistical differentiation, centroids occupied distinct positions in the nMDS environmental space (Fig. 2). Reproductive adults exhibited the most negative centroid shift along both axes (nMDS1 = -0.08, nMDS2 = -0.15), while seedlings showed the greatest positive shift along nMDS1 (0.55) and saplings along nMDS2 (0.51) axes. Non-reproductive adults occupied an intermediate position but showed the highest dispersion (nMDS1 SD = 1.64). The largest Euclidean distance between centroids was observed between seedlings and reproductive adults (0.72 units), followed by the separation between saplings and reproductive adults (0.69 units). The distance between seedlings and saplings was moderate (0.54 units), while non-reproductive adults showed lower separation from reproductive adults (0.53 units).

**Figure 2.**
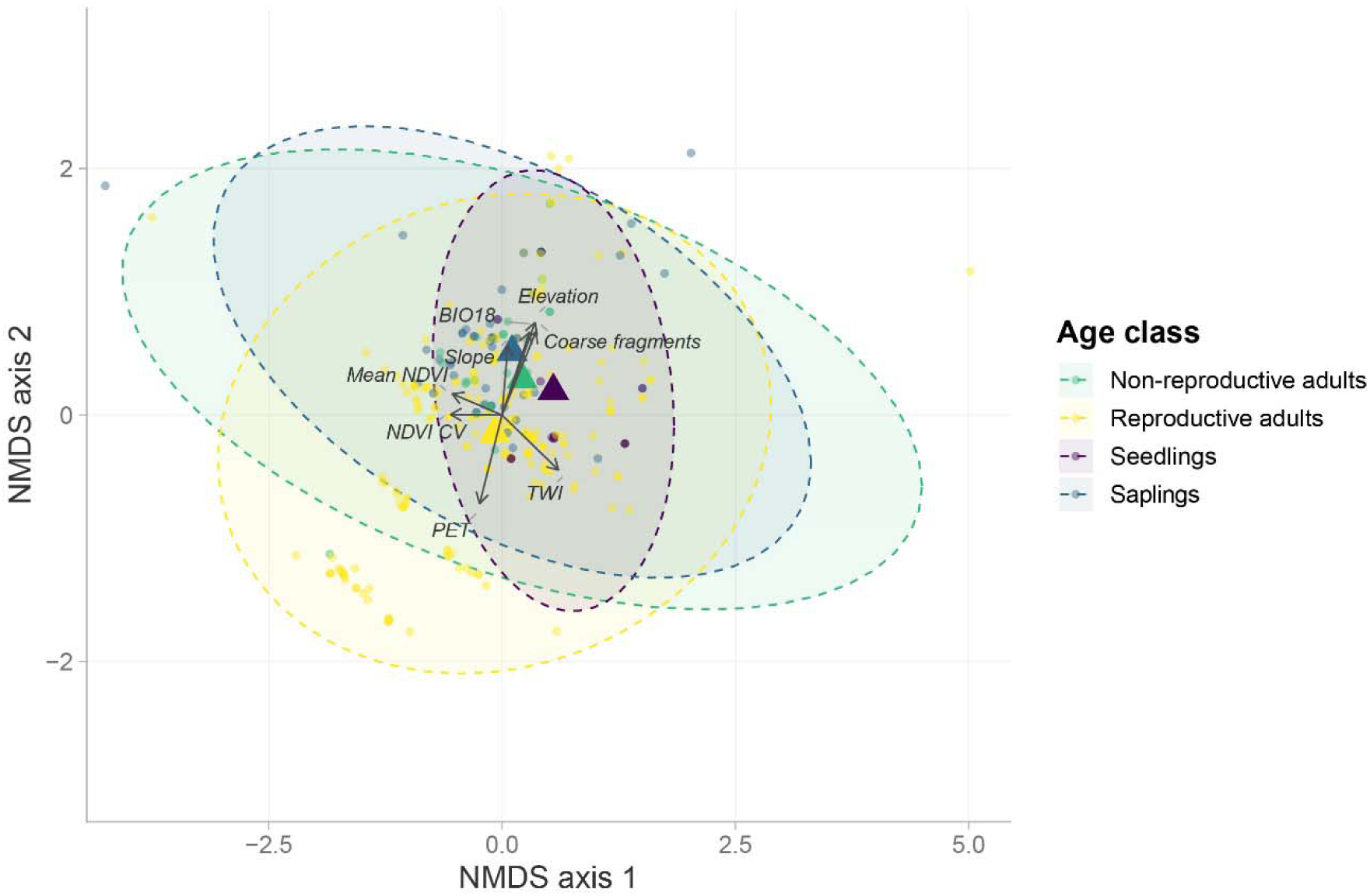
Non-metric multidimensional scaling (nMDS) ordination of standardized environmental conditions occupied by age classes of *Jubaea chilensis* (Stress = 0.15; PERMANOVA: F_3,328_ = 4.51, p < 0.0001). Distances between points approximate environmental dissimilarity, with closer points indicating more similar niches. Points represent individual plants; triangles indicate class centroids; ellipses delimit 95% confidence regions. Arrows show environmental variables significantly associated with the ordination (vector fitting, R² > 0.3, p < 0.05), with arrow direction indicating the gradient and arrow length proportional to the strength of the association (R²).

Eight environmental variables showed strong association and statistical significance with the nMDS ordination (R² > 0.30, p ≤ 0.001). Elevation was identified as the most strongly correlated variable with the ordination, explaining 69.7% of the variation in the NMDS space (R² = 0.70, p = 0.001), followed by variables related to water balance: coarse fragments (R² = 0.60), PET (R² = 0.58), TWI (R² = 0.57), and precipitation of the warmest quarter (R² = 0.54). Furthermore, topographic and vegetation variables also showed significant, although moderate, relationships: slope (R² = 0.32), mean NDVI (R² = 0.32), and NDVI CV (R² = 0.31).

### Environmental filters on recruitment

The multiclass Random Forest model showed good overall performance (OOB error = 24.1%), with marked differences in predictive accuracy among age classes, consistent with environmental differentiation across ontogenetic stage. Variable importance based on Mean Decrease Gini was dominated by vegetation-related predictors, led by mean NDVI (28.10), followed by NDVI temporal trend (20.50), NDVI seasonal amplitude (19.10), NDVI sensitivity to ENSO (18.50), and TWI (18.50) (Fig. 3a). On the other hand, binary Random Forest showed an OOB error of 15.7%, with sensitivity (ability to detect true seedlings) of 80% and specificity (ability to identify non-seedlings) of 84.5%. Mean Decrease Accuracy importance ranking showed a different set of predictors compared with the multiclass model. NDVI CV was identified as the dominant predictor with MDA = 4.93, followed by orientation along the east-west axis (eastness) with MDA = 3.64, and texture index with MDA = 3.01. Cation exchange capacity and NDVI seasonal amplitude were also ranked among the dominant predictors, although with less importance than the first three (MDA = 0.17 and 0.04, respectively) (Fig. 3b). Additionally, the binary Random Forest model trained to discriminate between saplings and reproductive adults showed an accuracy slightly superior to the seedling model (OOB error = 13.7%). Importance ranking revealed a different set of predictors, with elevation being the most important one (MDA = 32.50), followed by slope (MDA = 19.90), mean NDVI (MDA = 19.60), TWI (MDA = 19.50), and NDVI temporal trend (MDA = 18.20). Overall, seedling recruitment is constrained by microhabitat conditions, whereas saplings and reproductive adults are differentiated primarily by topography.

**Figure 3.**
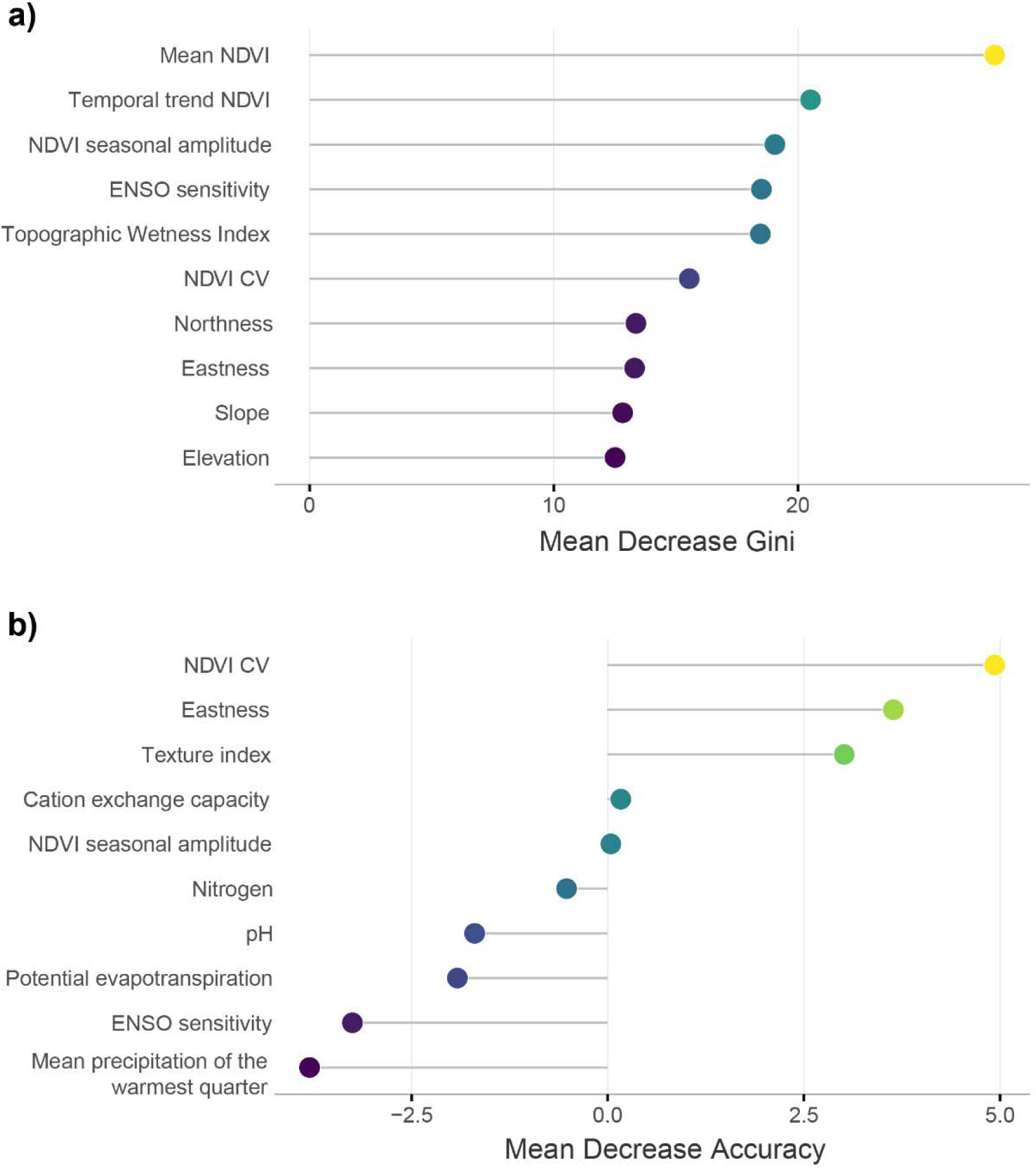
Variable importance from Random Forest models predicting ontogenetic stage in *Jubaea chilensis*. a) Mean Decrease Gini for the multiclass model (seedlings, saplings, non-reproductive adults, reproductive adults; OOB error = 24.1%); top 10 of 18 predictors shown. b) Mean Decrease Accuracy (MDA) for the binary seedling vs. non-seedling model (OOB error = 15.7%; sensitivity = 80%; specificity = 84.5%); top 10 of 18 predictors shown. Near-zero or negative MDA values indicate negligible contribution to seedling classification.

### Spatial distribution of suitability and niche contraction

Predicted probabilities derived from the binary Random Forest model clearly differentiated age classes. Seedlings showed high predicted probabilities (median = 0.96), whereas all remaining classes exhibited lower and broadly overlapping values (medians ranging from 0.11 to 0.19). The comparison between seedlings and non-seedlings showed significant differences (Wilcoxon test: p < 0.001), indicating that the model consistently discriminates the environmental conditions associated with successful seedling establishment. Notably, despite most reproductive adults showing low predicted suitability (50% with probability < 0.11), some individuals exhibited high probabilities (maximum = 0.95), making them indistinguishable from seedlings based on predicted suitability alone, and indicating environmental overlap between historical and current occupancy conditions.

On the other hand, the continuous map revealed marked spatial heterogeneity in predicted recruitment suitability (Fig. 4). Spatial structure was non-random, with patches of high probability (probability > 0.60) alternating with zones of low suitability (probability < 0.20). The classification of all individuals based on their predicted suitability and actual class showed an important decoupling between environmental suitability and realized recruitment. Seedlings represented only 4.5% of the total, corresponding to sites where recruitment was successful. Importantly, 21.1% of individuals were classified as high suitability without seedlings, representing sites where environmental variables predict suitable conditions, but recruitment has not occurred. Furthermore, 24.4% of individuals occupied sites of medium suitability, and 50% occupied sites of low suitability. Class composition of sites with failed recruitment revealed that most individuals belonged to reproductive adults (68.6%), followed by saplings (22.9%) and non-reproductive adults (8.6%).

**Figure 4.**
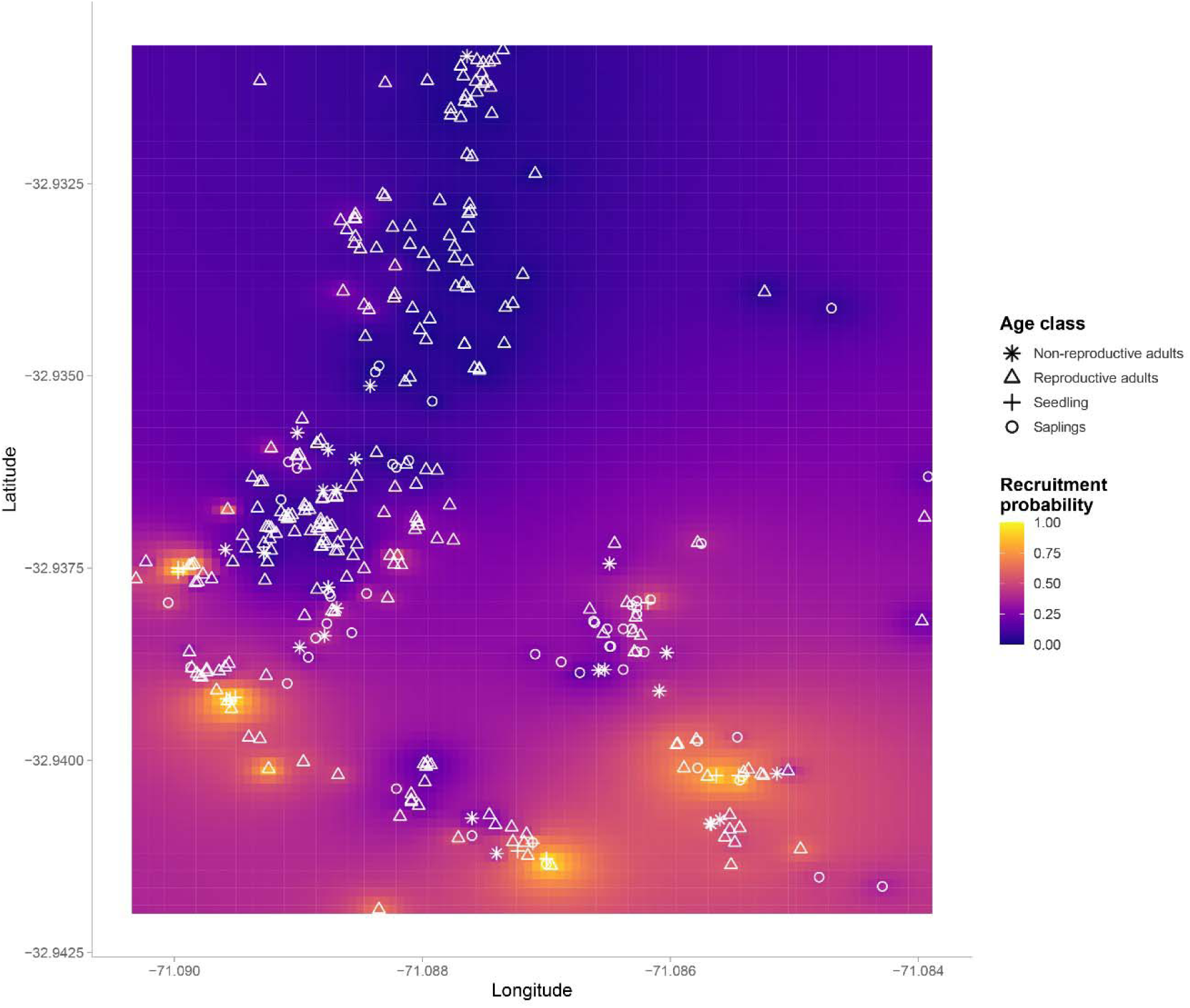
Spatial distribution of *Jubaea chilensis* individuals overlaid on predicted recruitment probability, estimated by inverse distance weighting interpolation of Random Forest predictions of seedling establishment. Areas of high predicted probability (yellow) occupied predominantly by non-seedling individuals indicate sites of apparent recruitment failure, where environmental conditions are suitable but recruitment has not occurred.

Lastly, a comparison between the adult niche and the seedling niche for the five most important predictors identified by the seedling Random Forest model showed consistent directional shifts, although none were statistically significant (all Wilcoxon p > 0.10). The largest relative change was observed for NDVI CV (-44%), followed by a shift in east-west orientation, suggesting a tendency toward contraction of the regeneration niche, although this pattern is not statistically supported.

## Discussion

Our results are consistent with environmental filtering throughout ontogeny, combined with the decoupling between predicted suitability and realized recruitment, as drivers of the regeneration collapse of *J. chilensis*. Nevertheless, experimental approaches such as seed and seedling transplants across the suitability gradient would be needed to confirm this further. The spatial segregation among age classes indicates a shift in the spatial scale of environmental filtering across ontogeny. Seedling establishment is primarily associated with microhabitat conditions, whereas the persistence of reproductive adults is primarily associated with landscape-scale topography. This scale-dependent mismatch decouples the presence of adult individuals from environmental suitability for recruitment. The significant spatial aggregation observed, with decreasing intensities from seedlings to reproductive adults, is consistent with the patterns reported in Amazonian long-lived palm species (Choo et al. 2017). This pattern also aligns, to some degree, with the Janzen-Connell hypothesis: most seeds fall within the immediate vicinity of parental plants, increasing density-dependent mortality due to seed predation and intraspecific competition, favoring individuals dispersing to longer distances (Connell 1971, Janzen 1971). Since *O. degus* disperses seeds over short distances (mean 6.2 ± 0.4 m; Cordero et al. 2021), spatial clustering of seedlings likely reflects both this limited dispersal range and subsequent environmental filtering of caches, rather than dispersal behavior alone. However, the observed segregation is not limited to a gradient of parental distance but reflects a differential occupation of environmental space (Barton 2024). Five of six pairwise comparisons between age classes were significant, with the greatest distance in environmental space observed between seedlings and reproductive adults, suggesting that spatial segregation is a consequence of the contrasting environmental conditions required by each ontogenetic stage.

Although environmental differentiation among age classes in palm species has been previously documented (Choo et al. 2017, Oda et al. 2019), the environmental drivers underlying this differentiation across the complete ontogenetic sequence remain scarcely understood. Our results show that the differentiation between reproductive adults and saplings is dominated by topography at a landscape scale, while the establishment of seedlings depends mainly on microhabitat. This change in environmental predictors among ontogenetic stages is consistent with the idea that the fundamental niche of a species is not static over the life cycle but reconfigures based on changes in size, physiology and capacity to tolerate environmental stress (Bertrand et al. 2011, Barton 2024). In large palm species like *J. chilensis*, this transition is especially marked since adults with deep root systems and large water reserves in the trunk can thrive under xeric conditions and topographically exposed sites (González et al. 2017). Conversely, seedlings are completely dependent on superficial soil moisture and understory shade, being unable to establish without favorable conditions (Guzmán et al. 2017).

We observed that NDVI CV is the most dominant predictor of seedling recruitment, which suggests high temporal variability in vegetation cover reflecting sites where the cover is neither homogeneous nor constant, such as scrub edges, small clearings, or transition zones and open scrublands. These microsites with vegetation heterogeneity offer a combination of moderate solar radiation, partial cover from summer drought, and less competition for resources from the closed canopy (Gómez-Aparicio et al. 2005, Lloret et al. 2005). Noteworthy, these microsites are also preferred by *Octodon degus*, the only seed disperser of *J. chilensis* seeds (Cordero et al. 2021, Cordero et al. 2026), which adds another layer of complexity to palm recruitment, determined by biotic interactions as discussed earlier. Moreover, eastness also plays an important role, suggesting that seedling tend to establish on hillsides oriented to the east, which receive morning radiation in the southern hemisphere but are protected from direct radiation during the afternoon. Since the latter is the most intense during the prolonged Mediterranean summer, it allows reducing evapotranspiration in seedlings with limited root systems (Badano et al. 2005). Soil texture also plays a critical role, with intermediate or slightly fine texture soils retaining more moisture available for germination (Muñoz-Rojas et al. 2016), in contrast with coarse texture which is predominant in zones of higher elevation and slope (Tesfa et al. 2009), where reproductive adults are more frequent.

On the other hand, environmental predictors differentiating saplings from reproductive adults mainly belong to topographic gradients at a landscape scale and mean vegetation productivity. This suggests that individuals persisting past the critical establishment stage are progressively more to be found in microsites less constrained by microenvironmental conditions, resulting in an environmental space more similar to that of adults. It is important to note, however, that age classes here correspond to different individuals compared at a single point in time, not the same plants followed through their lifespan. Therefore, this topographic segregation should not be understood as an ontogenetic shift within an individual (a sessile plant cannot move to a different slope or elevation). A more feasible explanation is differential survival among individuals that established decades or centuries apart. The topographic positions where reproductive adults now stand may have been suitable microsites at the time those individuals germinated, under vegetation cover, climate, or disturbance conditions quite different from those shaping recruitment today.

Notably, 21.1% of the sampled individuals occupy sites with high predicted environmental suitability for seedling recruitment, without this recruitment taking place. Inversely, only 4.5% of individuals were observed as seedlings in the field. Although recruitment in long-lived species can occur in irregular pulses, recruitment failure in this system has been documented through the full seed dispersal process, with an estimated 1.81% seed-to-seedling success rate (Cordero et al. 2021). This decoupling suggests that suitable microsites for establishment exist across the landscape, yet recruitment fails to occur in most of them. The sites of high suitability without seedlings are mainly dominated by reproductive adults (68.6%) and saplings (22.9%), consistent with older individuals persisting in microsites without new recruitment. This phenomenon may be explained by the low efficiency of the seed disperser, the native rodent *O. degus*, which preys on a significantly higher number of seeds than the ones it disperses, allowing only a small fraction of seeds to be available for germination (Cordero et al. 2021). Therefore, even though the environmental suitability of microsites is high and the disperser is present, this decoupling is consistent with insufficient effective dispersal, limiting the translation of environmental suitability into realized establishment (Turnbull et al. 2000, Muller-Landau et al. 2002).

The comparison between the environmental conditions occupied by reproductive individuals (adult niche, established at least ∼60 years ago) and those occupied by seedlings (seedling niche) did not reveal statistically significant differences in any of the five most important variables identified by the binary model, the multivariate analysis, or the permutation tests by variable. However, this result should be interpreted with caution, as the sample size of seedlings (n = 15) lacks statistical power to detect real differences, particularly given the highly asymmetric sample sizes between seedlings and reproductive adults (n = 242). Therefore, only effects of large magnitude would be detectable, and the absence of significance is not equivalent to absence of effect. Nevertheless, the observed tendency in some variables is coherent with predictions derived from the regional environmental context. The highest relative change in medians was observed in NDVI CV, suggesting that current seedlings tend to establish at microsites with lower temporal variability in vegetation cover than those occupied by reproductive adults when they established. The east-west orientation showed a similar direction of change, although also lacking statistical support. These trends, however, are only suggestive of regenerative niche contraction under Mediterranean aridification, as this hypothesis was not confirmed with the present data. During more than a decade, the megadrought in central Chile reduced seasonal water availability (Boisier et al. 2016, Garreaud et al. 2020), potentially restricting successful recruitment to an increasingly narrow set of microsites.

However, this interpretation cannot be completely separated from purely ontogenetic differentiation in environmental tolerances between seedlings and adults, which would arise regardless of any climatic shift. It is also difficult to rule out possible rapid evolutionary change in the genotypic composition of recruiting seedlings relative to established adults. Disentangling these alternatives, and testing whether recruitment is tied to ephemeral, interannual pulses in water availability rather than long-term mean conditions, would require long-term demographic monitoring or common-garden approaches beyond the scope of this study. As noted above, this cross-sectional comparison of different individuals is a proxy for, and not a direct measure of, the demographic vital rates underlying recruitment. This contrasts with the recruitment failure itself, which has independent temporal support from historical census comparisons and seed-to-seedling tracking(Bravo et al. 2018, Cordero et al. 2021). Furthermore, this study characterizes a single population, and replicate sites spanning a gradient of disturbance would provide a more robust evaluation of how representative these patterns are of the species’ broader, more fragmented range. However, the comparably sized population (El Salto palm grove) currently shows no recruitment (Cordero et al. 2021), precluding a direct replicate comparison at present.

The spatial contraction of the geographic area occupied by seedlings relative to that occupied by reproductive adults indirectly supports this argument. Regardless of whether the environmental conditions of microsites have changed, the reduction in the effective recruitment area implies that fewer individuals have access to suitable sites for establishment. Considering the already low recruitment rate (∼1.81%; Cordero et al. 2021), this represents a clear extinction debt, masked only by the longevity of *J. chilensis*. Similar patterns have been observed in the palm *Euterpe edulis* in the Brazilian Atlantic Forest, where fragmentation and aridification severely compromised recruitment with long-term consequences for population viability (Pizo et al. 2006). However, the Mediterranean distribution of *J. chilensis* intensifies the problem, as recruitment is restricted to narrow temporal windows associated with winter rainfall events (Holmgren et al. 2006).

*Jubaea chilensis* is a narrowly distributed, monotypic relict, but its restricted range does not appear to reflect a narrow physiological niche. The species tolerates temperatures as low as -22°C in cultivation and thrives well beyond its natural range under milder Mediterranean and even temperate climates (Guzmán et al. 2017). Its current distribution reflects a drastic population contraction, from an estimated 5,000,000 individuals five centuries ago to around 123,000 today, compounded by historical isolation from tropical relatives across the Atacama’s Arid Diagonal (Guzmán et al. 2017). This limits the generalizability of our findings, as the ontogenetic recruitment failure we document may be relevant to other long-lived species undergoing range contraction, but *J. chilensis* also depends on a sole seed disperser (*O. degus*), whose role is increasingly compromised by an invasive species (Cordero et al. 2021). This dependence on a sole seed disperser is a structural vulnerability more characteristic of narrowly distributed relict species than of widely distributed congeners with more redundant disperser assemblages.

## Conclusion

Our results have direct implications for *J. chilensis* management. Microsites with high suitability for recruitment should be considered priority candidates for active interventions through direct seeding or assisted transplanting under suitable environmental conditions. Conservation strategies limited to protecting reproductive adults are insufficient, since our results suggest that adult presence is an unreliable indicator of suitable conditions for recruitment. In this sense, models based on adult distribution overestimate the amount of habitat available for effective recruitment and underestimate the urgency of the demographic situation. The protection of microsites with environmental suitability for seedlings, including the reduction of predation pressure, represents the most direct strategy to increase the recruitment rate above the current critical threshold. Future management programs must simultaneously address the environmental and biotic filters of recruitment; failing to do so risks making local extinction inevitable within the next century.

